# Construction of an index system for big data application in health care enterprises --A study based on the Delphi method

**DOI:** 10.1101/477067

**Authors:** Yue Chang, Wei Du, Ruo Shi, Tang Lei, Hanni Zhou, Duan Li, Jiaqi Hu, Xingyin Fan

## Abstract

This study explored the theory of medical enterprise management and big data. Based on the Delphi method, two rounds of expert opinions were consulted on the capability of a health care enterprise big data application index system covering 11 dimensions, 46 first-level indicators and 111 second-level indicators. The index system includes two categories of input and capacity. The input category includes five dimensions: human resources, material resources, financial resources, government policies, and social service system or social environment. The capabilities category includes six dimensions: data integration capabilities, service capabilities, data analysis capabilities, information security, profitability, and innovation capabilities. This index system aims to appraise the application capability of big data scientifically and systematically, and fills the gaps of such research in China so far, which has positive significance for further research in the future.

## Introduction

*The Big Health Industry of China* consists of two services: medical health services and non-medical health services. The industry has formed four basic industrial groups, namely *Medicine*, *Wellness*, *Health*, and *Management*. Among these, the health-raising industry, supported by *Wellness*, covers the whole life cycle, surrounding human body and mind, integrating medical services, big data health information services, health management and promotion services, health insurance services and other ancillary services. *The Wellness industry of China* consists of four major formats: *Leisure Wellness*, *Nourishing Wellness*, *Sports Wellness*, and *Hot Spring Wellness*. In 2017, Xi Jinping, Chinese President, pointed out in the report of the 19^th^ National Congress that "during the implementation of the healthy China strategy, the people's health is an important symbol of national prosperity. We must improve the national health policy and provide all-round full-cycle health services for the people." It is predicted that the health industry will lead a new round of development boom in China, then spread to the whole world.

With the continuous popularization of network and information technology, the amount of data generated by humans is growing exponentially. Now we are entering the era of big data. The application of big data has penetrated into every industry and business and has gradually become an important basis for predicting industry markets, making decisions, and understanding competitors. For health care companies, big data applications will change their operating models and management methods. The application of big data to health care enterprises with rich connotations is a relatively new topic. The wave of big data is not only a revolution in the field of information technology, but also a tool to accelerate marketing changes and lead social change on a global scale[1].

In stark contrast to the rapid development of big data in all walks of life, there is currently no research in the world on the construction of an index system for big data and its applicability for health care enterprises. In May 2001, McKinsey first mentioned *big data* in a research report titled *Big Data: The Next Frontier of Innovation, Competition, and Productivity.* Before 2013, big data was still a new concept, which was introduced in few related studies, small number of introductory, predictive articles. Today, just a few years later, we discovered that big data has great power to accurately reflect the real world and is of great value. Managers can make decisions based on big data, so that enterprises can get greater benefits on less investment basis. Therefore, it is imperative to develop such a system and scientifically evaluate health care enterprises’ big data application, and to provide supervision standards for government regulatory authorities. This paper focuses on the big data application capability, evaluating the results of the big data application for health care enterprises.

In the 19^th^ century, the French economist Jules Dupuit proposed a cost-benefit theory, which posits that the ultimate goal pursued by any enterprise is efficiency. In order to benefit, costs must be paid [2]. Dupuit argues that the implementation of any policy always requires a certain amount of human, material and financial resources, thus these resources cannot be used for other policies. This is the cost of a policy. The theory requires that enterprises should know the cost-effective concept and consider the necessity and rationality of *cost* by comparing *input* and *output*. The *United Nations Development Programme* proposes that capability is the competence and strength of an organization or individual to perform functions consistently, effectively and efficiently [3]. Based on these two important theories, this paper set two categories of the index system: costs and capabilities. Human, material, and financial resources are the three dimensions of the cost category.

Using big data, health care enterprises will collect many isomorphic and heterogeneous data, so noise data interference will appear in a database. Thus the enterprise needs to perform data integration such as data cleaning and data conversion on the original data[4]. The enterprise will then integrate data and form big data analytics to generate profitability for their business[5]. As a service-oriented industry, the enterprise must respond quickly to customer needs and improve customer satisfaction by using big data. Jin Hao (2003) proposed that, in a competitive market, a company must continue to provide products or services to the market more effectively than other companies[6] and gaining profit and self-development is an essential capability of any enterprise. In the study of enterprise input-output performance, the well-known Cobb-Douglas function, proposed by Charles Cobb and Paul Douglas, describes the value of an output brought about by innovation and is widely used in the macro-economic and micro-enterprise innovations. A report by James McKinsey also stresses the role of innovation in enterprise big data applications[7]. In summary, this paper sets the dimension of the capability category as data integration capability, data analysis capability, service capability, profitability, and innovation capability as the first round of questionnaires for expert consultation.

## Methods

### Literature Search

We collected research articles related to the health care industry and big data. Sources included domestic and foreign papers, dissertations, and books. Keyword search terms included *big data application*, *index system for big data application*, and *health care industry*. We then determined the overall framework of the index system in health care enterprises.

### Expert interview

We interviewed experts from the Guizhou Provincial Development and Reform Commission, Guizhou Provincial Health Planning Commission, Guiyang Big Data Development Management Committee, Guizhou University and Guizhou Medical University, and Guizhou Lianke Health Information Technology Co., Ltd. to understand the current status of big data applications and asked them for suggestions for the index for big data application in health care enterprises. Finally, we formulated a preliminary index system.

### Delphi method

After formulating the index system, we used the Delphi method to screen the indicators. The steps are as follows:

1. Setting up a research team: The research team consisted of five members who prepared questionnaires, selected and confirmed the experts, and analyzed the results.
2. Selecting the inquiry expert: We pre-selected 17 experts based on the following selection criteria: 1) government department managers with certain decision-making significance on the popularization and application of big data; 2) enterprise managers with rich experience in big data; 3) scholars engaged in big data-related research; 4) experts who are interested in this research; 5) medium grade professional title.
3. Designing a questionnaire.
4. Consulting with experts: The questionnaire was distributed to the experts in an anonymous or back-to-back manner. After two rounds of consultation and feedback, the index system was formed.
5. Statistical analysis: Data were computerized and analyzed using SPSS version 21.0.

### Indicator screening principle

1. Scientific principle: The indicator conforms to the criteria of objective facts and has a scientific theoretical basis. The definition of each indicator is clear and precise, and the specific calculation formula is expressed for indicators that are prone to ambiguity.
2. Comprehensive principle: Through systematic investigation and demonstration, all indicators that measure the big data application capability are covered as much as possible.
3. Principle of operability: The index system should reflect the direction of big data development and have guiding significance for health care enterprises to achieve further big data application. Therefore, the indicators should be universally defined. The meaning of the indicators should be observable, measurable, and operational.

### Indicator screening method

Each indicator was rated on a scale of 1 to 5 where 1 represents very unimportant and 5 represents very important. Scores for each dimension and indicator were summed. The experts’ evaluation of each indicator was measured using three statistical measures: the percentage of experts who held ‘very important’ and ‘important’ opinions, the mean score, and the coefficient of variation. An indicator was deleted if the percentage of experts that felt it was important was less than 75%, the mean score was less than 4.0, and the coefficient of variation was greater than 1.0. If only one or two of these statistical measures were met, the retention of the indicator was determined after discussion by the research team.

### Characteristics of Experts

This study pre-selected 17 experts and received feedback from 16. The 16 experts were senior teachers from various universities in Guizhou province, heads of big data companies, or persons from relevant departments. They all had systematic and unique insights in the area of big data. The basic information of the 16 experts is shown in Table 1.

**Table 1.** Expert characteristics.

| Category | Item | Number | Percentage |
| --- | --- | --- | --- |
| Gender | Male | 11 | 69 |
|  | Female | 5 | 31 |
| Educational background | Bachelor's degree | 2 | 13 |
|  | Master's degree | 5 | 31 |
|  | Doctoral degree | 9 | 56 |
| Title | Middle Title | 4 | 25 |
|  | Vice-senior Title | 9 | 56 |
|  | Senior Title | 3 | 19 |
| Occupational position | Management | 2 | 13 |
|  | Technology | 5 | 31 |
|  | Research | 9 | 56 |

## Results and discussion

### First round of inquiry

#### Questionnaire setting and distribution

The purpose of the first round of questionnaires was to ask experts to comment on the importance of the researcher's initial setting of the indicators for big data application capability. The questionnaire asked for evaluation of each dimension, the first-level indicators, and the second-level indicators in the index. In each part, the items of modification opinions and other indicators were set up, and experts were requested to propose amendments.

The first round of questionnaires was issued to 17 experts, of which 16 were recovered resulting in a recovery rate of 94%.

#### Reliability

Chronbach's alpha values of the whole system and for the first and second levels were 0.96, 0.85, and 0.96, respectively, indicating that the reliability of the index system was quite high.

#### Statistical Analysis

Table 2 shows the importance percentage for each dimension and indicator selected by the 16 experts, together with the mean, standard deviation, and coefficient of variation.

**Table 2.** Importance for dimensions

| Category | Dimension | Importance (%) |  |  |  |  | Mean | Standard deviation | Coefficient of variation |
| --- | --- | --- | --- | --- | --- | --- | --- | --- | --- |
|  |  | Very important | Important | General | Unimportant | Very unimportant |  |  |  |
| Costs | Human Resource | 69 | 31 | 0 | 0 | 0 | 4.69 | 0.48 | 0.10 |
|  | Material Resource | 25 | 56 | 19 | 0 | 0 | 4.06 | 0.68 | 0.17 |
|  | Financial Resource | 50 | 50 | 0 | 0 | 0 | 4.50 | 0.52 | 0.11 |
| Capabilities | Data Integration | 69 | 19 | 6 | 0 | 6 | 4.44 | 1.16 | 0.26 |
|  | Service | 25 | 56 | 19 | 0 | 0 | 4.06 | 0.68 | 0.17 |
|  | Data Analysis | 75 | 19 | 6 | 0 | 0 | 4.69 | 0.60 | 0.13 |
|  | Profitability | 19 | 75 | 6 | 0 | 0 | 4.13 | 0.54 | 0.13 |
|  | Creativity | 44 | 56 | 0 | 0 | 0 | 4.44 | 0.52 | 0.12 |

As shown in Table 2, the average score of the eight dimensions was above 4.0 (important), the important percentage was higher than 90%, and the coefficient of variation was less than 1.0 indicating that the experts believed these eight dimensions were highly and consistently important.

Table 3 shows that in the COSTS category, the mean of ‘Personnel’ and ‘Other Costs’ was less than 4.0. In the capabilities category, the mean of ‘Service Coverage’, ‘Government Big Data Subsidy Income’, and ‘Other Income’ were below 4.0. The percentage of experts who felt that these five indicators were important was less than 75. Apart from ‘3 Information Security’, the standard deviation of other first-level indicators was less than 1.0. The coefficient of variation for all indicators was less than 1.0.

**Table 3.** Importance for the first-level indicators

| Dimension | First-level Indicators | Importance (%) |  |  |  |  | Mean | Standard deviation | Coefficient of variation |
| --- | --- | --- | --- | --- | --- | --- | --- | --- | --- |
|  |  | Very important | Important | General | Unimportant | Very unimportant |  |  |  |
| Human Resource | 1. Personnel | 6 | 50 | 44 | 0 | 0 | 3.63 | 0.62 | 0.17 |
|  | 2. Qualification | 38 | 50 | 12 | 0 | 0 | 4.25 | 0.68 | 0.16 |
|  | 3. Staff Training | 56 | 38 | 6 | 0 | 0 | 4.50 | 0.63 | 0.14 |
| Material Resource | 4. Hardware | 38 | 44 | 19 | 00 | 0 | 4.19 | 0.75 | 0.18 |
|  | 5. Software | 63 | 31 | 6 | 00 | 0 | 4.56 | 0.63 | 0.14 |
| Financial Resource | 6. Assets Size and Proportion | 25 | 63 | 6 | 6 | 0 | 4.06 | 0.77 | 0.19 |
|  | 7. Operating Costs | 13 | 75 | 12 | 0 | 0 | 4.00 | 0.52 | 0.13 |
|  | 8. Service Costs | 13 | 81 | 6 | 0 | 0 | 4.06 | 0.44 | 0.11 |
|  | 9. Technical Upgrade Costs | 31 | 56 | 13 | 0 | 0 | 4.19 | 0.66 | 0.16 |
|  | 10. Technical Staff Costs | 69 | 31 | 0 | 0 | 0 | 4.69 | 0.50 | 0.11 |
|  | 11. Other Costs | 0 | 19 | 69 | 0 | 12 | 3.36 | 0.97 | 0.29 |
| Data Integration | 1. Collection | 63 | 37 | 0 | 0 | 0 | 4.63 | 0.50 | 0.11 |
|  | 2. Extracting Valid Data | 88 | 12 | 0 | 0 | 0 | 4.88 | 0.34 | 0.07 |
|  | 3. Information Security | 50 | 44 | 0 | 0 | 6 | 4.31 | 1.01 | 0.24 |
| Service | 4. Service Coverage | 6 | 63 | 31 | 0 | 0 | 3.75 | 0.58 | 0.15 |
|  | 5. Service Satisfaction | 31 | 63 | 6 | 0 | 0 | 4.25 | 0.58 | 0.14 |
| Data Analysis | 6. Analysis of Unstructured Data | 50 | 31 | 19 | 0 | 0 | 4.31 | 0.79 | 0.18 |
|  | 7. Real-time Insight Warning | 69 | 25 | 6 | 0 | 0 | 4.63 | 0.62 | 0.13 |
|  | 8. Precision Marketing | 56 | 44 | 0 | 0 | 0 | 4.87 | 0.51 | 0.11 |
|  | 9. Cost Reduction and Efficiency | 50 | 44 | 6 | 0 | 0 | 4.73 | 0.63 | 0.13 |
|  | 10. Personalized Customer Management | 44 | 50 | 6 | 0 | 0 | 4.38 | 0.63 | 0.14 |
| Profitability | 11. Operating Income | 25 | 75 | 0 | 0 | 0 | 4.25 | 0.48 | 0.11 |
|  | 12. Government Big Data Subsidy Income | 0 | 44 | 50 | 6 | 00 | 3.38 | 0.62 | 0.18 |
|  | 13. Other Income | 0 | 19 | 69 | 6 | 6 | 3.20 | 0.73 | 0.23 |
| Creativity | 14. Product Development | 56 | 44 | 0 | 0 | 0 | 4.56 | 0.51 | 0.11 |
|  | 15. Market Competitiveness | 31 | 63 | 6 | 0 | 0 | 4.25 | 0.58 | 0.14 |
|  | 16. Technology Innovation Management | 38 | 62 | 0 | 0 | 0 | 4.38 | 0.60 | 0.14 |

Table 4 shows the experts’ feedback of the second-level indicators. In the Costs category, the following indicators had importance percentages less than 75%: ‘1.1 Number of Technicians’, ‘1.3 Number of Operators’, ‘2.1 Educational Background’, ‘2.3 License’, ‘3.1 Number of Internal Training per capita in past 3 years’, ‘3.2 Number of External Training per capita in past 3 years’, ‘4.3 Network Equipment’, ‘4.4 Storage Device’, ‘6.1 Total Assets’, ‘10.1 Proportion of Technical Staff Salary and Training Costs’, ‘11.1 Proportion of Other Costs’. The mean scores for ‘1.1 Number of Technicians’, ‘1.3 Number of Operators’, ‘2.1 Educational Background’, ‘2.3 License’, ‘3.1 Number of Internal Training per capita in past 3 years’, ‘3.2 Number of External Training per capita in past 3 years’, ‘4.3 Network Equipment’, ‘6.1 Total Assets’, ‘11.1 Proportion of Other Costs’, ‘6.2 The ratio of big data hardware and software assets to total assets’, ‘8.1 Proportion of Service Costs’ were less than 4.0. The standard deviation of ‘2.3 License’, ‘4.2 Data Center or Engine Room’, ‘6.2 The ratio of big data hardware and software assets to total assets’ were greater than or equal to 1.0. The coefficient of variation for all indicators was less than 1.0.

**Table 4.** Importance for the second-level indicators

| First-level Indicator | Second-level Indicator | Importance ( %) |  |  |  |  | Mean | Standard deviation | Coefficient of variation |
| --- | --- | --- | --- | --- | --- | --- | --- | --- | --- |
|  |  | Very important | Important | General | Unimportant | Very unimportant |  |  |  |
| 1. Personnel | 1.1 Number of Technicians | 25 | 38 | 3 | 0 | 0 | 3.88 | 0.77 | 0.20 |
|  | 1.2 Number of Data Analysis Technicians | 69 | 19 | 12 | 0 | 0 | 4.56 | 0.73 | 0.16 |
|  | 1.3 Number of Operators | 6 | 50 | 38 | 6 | 0 | 3.56 | 0.73 | 0.20 |
| 2. Qualification | 2.1 Educational Background | 6 | 31 | 56 | 6 | 0 | 3.38 | 0.72 | 0.21 |
|  | 2.2 Work Experience | 63 | 31 | 6 | 0 | 0 | 4.56 | 0.63 | 0.14 |
|  | 2.3 License | 13 | 25 | 44 | 6 | 13 | 3.19 | 1.17 | 0.37 |
| 3. Staff Training | 3.1 Number of Internal Training per capita in past 3 years | 0 | 56 | 38 | 6 | 0 | 3.50 | 0.72 | 0.21 |
|  | 3.2 Number of External Training per capita in past 3 years | 19 | 31 | 44 | 6 | 0 | 3.63 | 0.89 | 0.24 |
| 4. Hardware | 4.1 Server | 8 | 37 | 19 | 6 | 0 | 4.06 | 0.93 | 0.23 |
|  | 4.2 Data Center or Engine Room | 44 | 38 | 6 | 12 | 0 | 4.13 | 1.02 | 0.25 |
|  | 4.3 Network Equipment | 31 | 38 | 25 | 6 | 0 | 3.94 | 0.93 | 0.24 |
|  | 4.4 Storage Device | 38 | 31 | 25 | 6 | 0 | 4.00 | 0.97 | 0.24 |
| 5. Software | 5.1 Data Integration Software | 38 | 44 | 19 | 0 | 0 | 4.19 | 0.75 | 0.18 |
|  | 5.2 Data Storage Software | 44 | 44 | 13 | 0 | 0 | 4.31 | 0.70 | 0.16 |
|  | 5.3 Data Cleaning Software | 56 | 38 | 6 | 0 | 0 | 4.50 | 0.63 | 0.14 |
|  | 5.4 Data Analysis Software | 69 | 31 | 0 | 0 | 0 | 4.69 | 0.48 | 0.10 |
| 6. Assets Size and Proportion | 6.1 Total Assets | 6 | 63 | 31 | 0 | 0 | 3.75 | 0.58 | 0.15 |
|  | 6.2 The ratio of big data hardware and software assets to total assets | 31 | 44 | 13 | 13 | 0 | 3.94 | 1.00 | 0.25 |
| 7. Operating Costs | 7.1 Proportion of Operating Costs | 13 | 75 | 13 | 0 | 0 | 4.00 | 0.52 | 0.13 |
| 8. Service Costs | 8.1 Proportion of Service Costs | 6 | 69 | 25 | 0 | 0 | 3.81 | 0.54 | 0.14 |
| 9. Technical Upgrade Costs | 9.1 Proportion of Technical Upgrade Costs | 19 | 69 | 13 | 0 | 0 | 4.06 | 0.57 | 0.14 |
| 10. Technical Staff Costs | 10.1 Proportion of Technical Staff Salary and Training Costs | 25 | 56 | 19 | 0 | 0 | 4.06 | 0.68 | 0.17 |
| 11. Other Costs | 11.1 Proportion of Other Costs | 0 | 25 | 63 | 13 | 0 | 3.13 | 0.62 | 0.20 |
| 1. Collection | 1.1 Timeliness of Collecting Data | 63 | 38 | 0 | 0 | 0 | 4.63 | 0.50 | 0.11 |
| 2. Extracting Valid Data | 2.1 Information System Connectivity | 38 | 56 | 6 | 0 | 0 | 4.31 | 0.68 | 0.16 |
| 3. Information Security | 3.1 Information Security Qualification | 31 | 56 | 13 | 0 | 0 | 4.19 | 0.72 | 0.17 |
|  | 3.2 Security Equipment | 44 | 44 | 6 | 6 | 0 | 4.25 | 0.91 | 0.21 |
| 4. Service Coverage | 4.1 Cumulative Number of Customer(individual/group) | 25 | 69 | 6 | 0 | 0 | 4.19 | 0.58 | 0.14 |
|  | 4.2 Service Radius | 13 | 38 | 50 | 0 | 0 | 3.63 | 0.79 | 0.22 |
| 5. Service Satisfaction | 5.1 Customers' Secondary Purchase Ratio | 13 | 88 | 0 | 0 | 0 | 4.13 | 0.40 | 0.10 |
|  | 5.2 Cycle of Repurchase | 25 | 63 | 6 | 6 | 0 | 4.06 | 0.81 | 0.20 |
|  | 5.3 Customer Satisfaction | 50 | 50 | 0 | 0 | 0 | 4.50 | 0.51 | 0.11 |
| 6. Analysis of Unstructured Data | 6.1 Proportion of Unstructured data | 13 | 56 | 25 | 6 | 0 | 3.75 | 0.77 | 0.21 |
|  | 6.2 Processing and Mining Unstructured Data | 50 | 50 | 0 | 0 | 0 | 4.50 | 0.50 | 0.11 |
| 7. Real-time Insight Warning | 7.1 Market Dynamic Monitoring Software | 38 | 44 | 19 | 0 | 0 | 4.19 | 0.75 | 0.18 |
|  | 7.2 Public Opinion Prediction Software | 38 | 50 | 13 | 0 | 0 | 4.25 | 0.66 | 0.15 |
| 8. Precision Marketing | 8.1 The number of People Captured Accurately | 25 | 63 | 13 | 0 | 0 | 4.13 | 0.62 | 0.15 |
|  | 8.2 Increasing of Conversion Rate | 31 | 69 | 0 | 0 | 0 | 4.31 | 0.46 | 0.11 |
|  | 8.3 Customer acquisition Costs Reduction | 31 | 69 | 0 | 0 | 0 | 4.31 | 0.48 | 0.11 |
|  | 8.4 Conversion Profits | 56 | 44 | 0 | 0 | 0 | 4.56 | 0.51 | 0.11 |
| 9. Cost Reduction and Efficiency | 9.1 Inventory Cost Reduction | 31 | 63 | 6 | 0 | 0 | 4.25 | 0.58 | 0.14 |
|  | 9.2 Labor Costs Reduction | 31 | 50 | 19 | 0 | 0 | 4.13 | 0.72 | 0.17 |
| 10. Personalized Customer Management | 10.1 Personalized Customer Management Capabilities | 31 | 56 | 13 | 0 | 0 | 4.19 | 0.62 | 0.15 |
|  | 10.2 Personalized Products and Offers* | 31 | 56 | 13 | 0 | 0 | 4.19 | 0.62 | 0.15 |
|  | 10.3 Customer Behavior Prediction Software | 43 | 57 | 0 | 0 | 0 | 4.43 | 0.51 | 0.12 |
| 11. Operating Income | 11.1 Operating Income | 44 | 50 | 6 | 0 | 0 | 4.38 | 0.62 | 0.14 |
| 12. Government Big Data Subsidy Income | 12.1 Government Big Data Subsidy Income | 6 | 38 | 44 | 13 | 0 | 3.38 | 0.81 | 0.24 |
| 13. Other Income | 13.1 Other Income | 0 | 33 | 60 | 7 | 0 | 3.27 | 0.59 | 0.18 |
| 14. Product Development | 14.1 Number of New Products Developed Each Year | 13 | 69 | 19 | 0 | 0 | 3.94 | 0.57 | 0.15 |
|  | 14.2 Number of New Technologies or Processes | 31 | 56 | 13 | 0 | 0 | 4.19 | 0.66 | 0.16 |
|  | 14.3 Response after products entering the market | 25 | 75 | 0 | 0 | 0 | 4.25 | 0.45 | 0.11 |
| 15. Market Competitiveness | 15.1 Whether The Latest Products Are Highly Differentiated | 38 | 50 | 13 | 0 | 0 | 4.25 | 0.68 | 0.16 |
|  | 15.2 Average Industry Market Share | 36 | 43 | 21 | 0 | 0 | 4.14 | 0.77 | 0.19 |
| 16. Technology Innovation Management | 16.1 Product Development and Innovation Management System | 38 | 63 | 0 | 0 | 0 | 4.38 | 0.50 | 0.11 |
|  | 16.2 Research & Development Center | 25 | 38 | 38 | 0 | 0 | 4.44 | 0.63 | 0.14 |

In the Capabilities category, the importance percentage of ‘4.2 Service Radius’, ‘6.1 Proportion of Unstructured data’, ‘7.1 Market Dynamic Monitoring Software’, ‘12.1 Government Big Data Subsidy Income’, ‘13.1 Other Income’, ‘16.2 Research & Development Center’ were below 75%. The mean scores of ‘4.2 Service Radius’, ‘6.1 Proportion of Unstructured data’, ‘12.1 Government Big Data Subsidy Income’, ‘13.1 Other Income’, ‘14.1 Number of New Products Developed Each Year’, ‘16.2 Research & Development Center’ were less than 4.0. The standard deviation and coefficient of variation for all indicators were less than 1.0.

#### Indicator screening results - first round

##### 1. Deleting

There were no indicators that met the criteria for deleting proposed by the research team, so no indicator was deleted.

##### 2. Adding

In the Costs category, one of the experts proposed to add a ‘Government Policy’ dimension and add five first-level indicators, namely ‘Technical Guidance’, ‘Funding Support’, ‘Policy Sustainability, ‘Strength’ and ‘Marketing Assistance’ and there corresponding secondary indicators. Another expert suggested adding a dimension of ‘Social Service System or Social Environment’, and adding two first-level indicators of ‘Social Information Infrastructure Construction’ and ‘Big Data Development Pilot Area’ and their corresponding secondary indicators.

In terms of the first-level indicators, one expert suggested adding ‘Synergy’ in the Human Resource dimension. ‘Synergy’ indicates the degree of collaboration within and outside the enterprise. In the ‘Data Integration’ Capability dimension, one expert recommended adding two first-level indicators, namely ‘Data Standards’ and ‘Heterogeneous System Data Integration’. In the ‘Service’ capability dimension, one expert suggested adding the first-level indicators of ‘Service Timely Response’. The discussion team determined the second-level indicators of ‘Service Timely Response Capability’ as ‘Customers’ Dependence on Products’, ‘Outlets’, ‘Service Speed’, ‘Service Quality’, ‘service quality tracking feedback’. In the ‘Data Analysis’ capability dimension, two experts recommend adding ‘Machine Learning’ and ‘Combination with Corresponding Business’ as two first-level indicators. In the dimension of ‘Creativity’, one expert suggested adding ‘Acceptance of New Ideas’ to emphasize the acceptance of new corporate ideas.

In terms of the second-level indicators, one expert suggested that ‘The Number of People with Work Experience in Well-known IT Companies (such as Baidu, Alibaba)’ is important for the application of big data. So it was added as the second-level of ‘Personnel’. One expert believed that ‘Security Equipment’ and ‘Database Middleware’ are important hardware for big data enterprises. In the ‘Financial Resource’ dimension, one expert proposed adding secondary indicators of ‘Core Technology Assets’. One expert held the opinion that ‘Information Security’ should be subdivided into two categories: ‘Enterprise Security’ and ‘Product Security’, adding corresponding second-level indicators. One expert suggested that ‘Artificial Intelligence’ and ‘Risk Control’ could be used to measure machine learning capability. Finally, one expert believed that obtaining financial support from national, provincial, municipal, district and county projects is important to the profitability of the company.

##### 3. Modifying

Two experts said that ‘Information Security’, which was originally a first-level indicator, was very important and should be a separate dimension. In terms of the first-level indicators, the original ‘Personnel’ was changed to ‘Staffing’, the original ‘Operating Costs’ was changed to ‘Operation and Maintenance Costs’, and the original ‘Collection’ was renamed to ‘Data Collection’, the original ‘Analysis of Unstructured Data’ was renamed to ‘Analysis of Structured or Unstructured Data’.

### Second round of inquiry

#### Questionnaire setting and distribution

According to experts' suggestions, the members of the research team decided to revise the indicators listed in the questionnaire and set up a second round of questionnaires. Significant changes included the addition of ‘Government Policy’ and ‘Social Service System or Social Environment’ dimensions in the input category, and the addition of an ‘Information Security’ dimension in the capacity category. Ma yanling summaried E. S. Masson, J. S. Bain, and Michael E. Poter’s competitive advantage external environment-based theory[8]. The investment of external factors to the enterprise is very important. The external policy environment, legal environment, and humanistic natural environment determine the number and distribution of the industry, determine market concentration and barriers to entry. In addition, one Chinese information expert (Qu Chengyi) raised vigilance and mentioned the importance of information security maintenance and supervision in big data management[9].

The results of the importance of indicators in second round were from 6 experts who gave feedback on the first round of questionnaires.

#### Reliability

The Chronbach Alpha values of the index system, the first level indicator, and the second level index were 0.98, 0.90, and 0.96 in the second round, indicating that the reliability of the index system was high.

#### Statistical Analysis

**Table 5.** Importance for dimension in the second round

| Category | Dimension | Importance (%) |  |  |  |  | Mean | Standard deviation | Coefficient of variation |
| --- | --- | --- | --- | --- | --- | --- | --- | --- | --- |
|  |  | Very important | Important | General | Unimportant | Very unimportant |  |  |  |
| Costs | Human Resource | 100 | 0 | 0 | 0 | 0 | 5.00 | 0 | 0 |
|  | Material Resource | 31 | 50 | 19 | 0 | 0 | 4.13 | 0.72 | 0.17 |
|  | Financial Resource | 56 | 44 | 0 | 0 | 0 | 4.56 | 0.51 | 0.11 |
|  | Government Policy | 38 | 56 | 6 | 0 | 0 | 4.31 | 0.60 | 0.14 |
|  | Social Service System or Social Environment | 6 | 88 | 6 | 0 | 0 | 4.00 | 0.37 | 0.09 |
| Capabilities | Data integration | 100 | 0 | 0 | 0 | 0 | 5.00 | 0 | 0 |
|  | Service | 25 | 63 | 13 | 0 | 0 | 4.13 | 0.62 | 0.15 |
|  | Data analysis | 94 | 6 | 0 | 0 | 0 | 4.94 | 0.25 | 0.05 |
|  | Information Security | 44 | 56 | 0 | 0 | 0 | 4.44 | 0.51 | 0.12 |
|  | Profitability | 19 | 56 | 25 | 0 | 0 | 3.94 | 0.68 | 0.17 |
|  | Creativity | 44 | 38 | 19 | 0 | 0 | 4.25 | 0.77 | 0.18 |

**Table 6.**
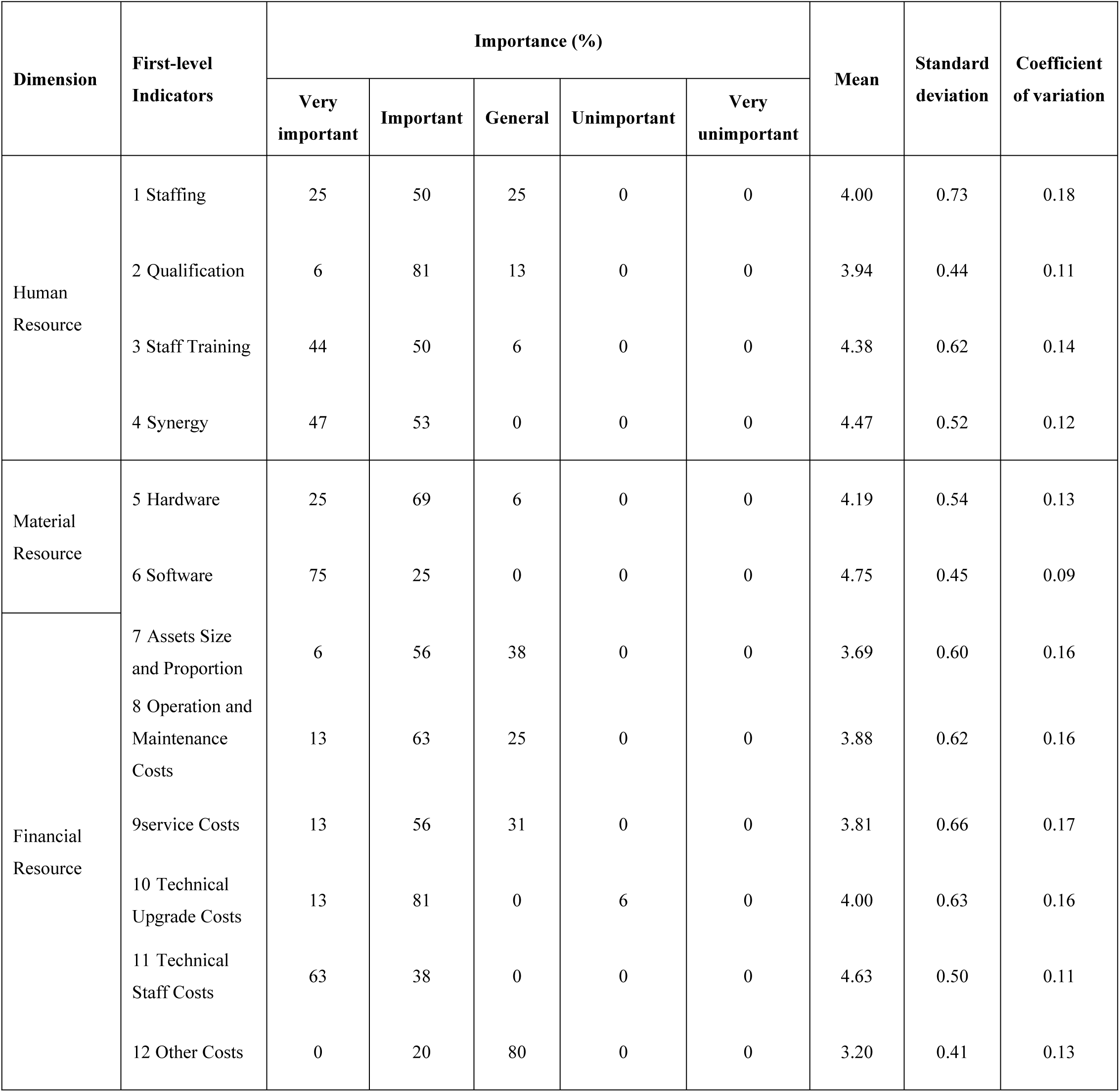
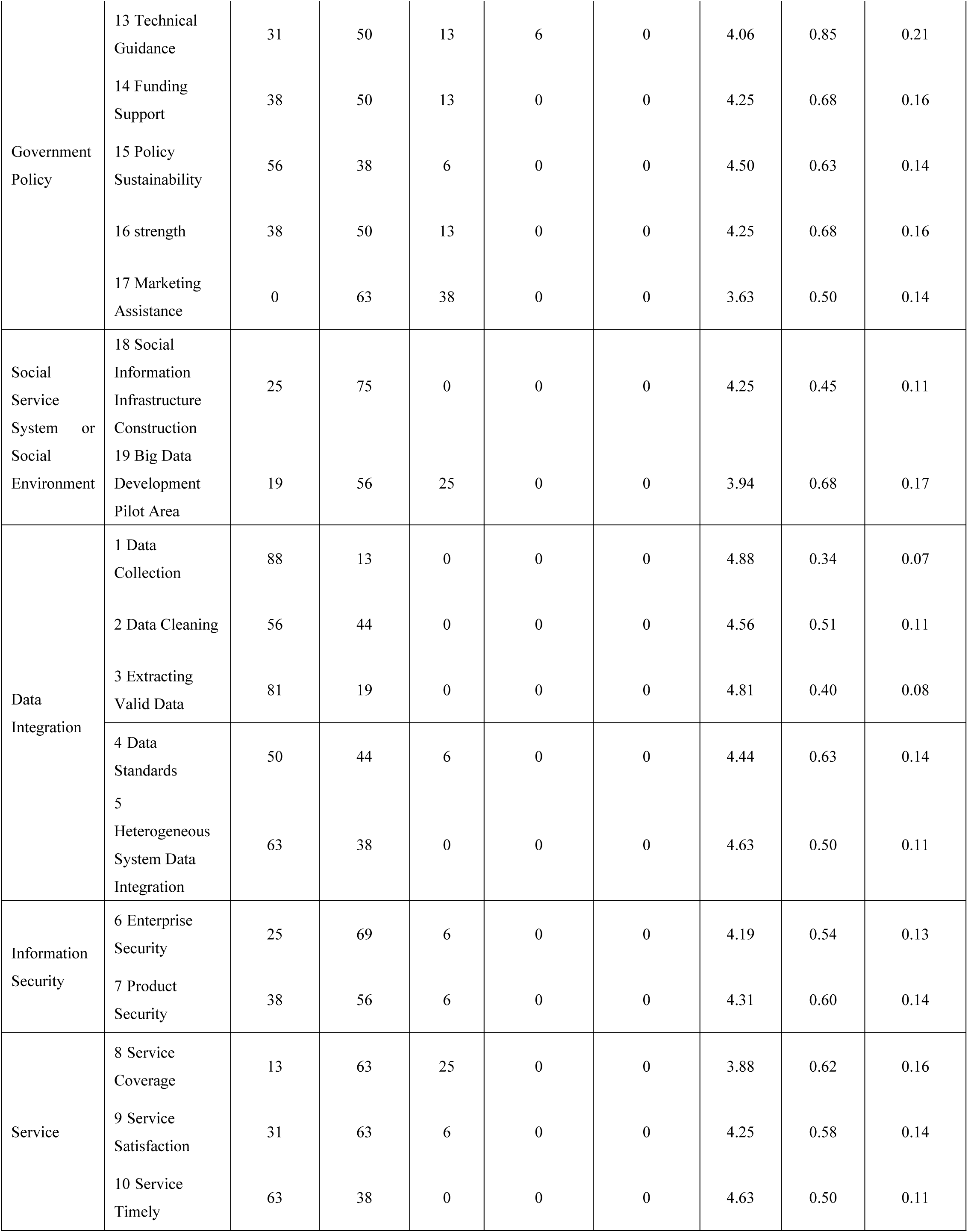
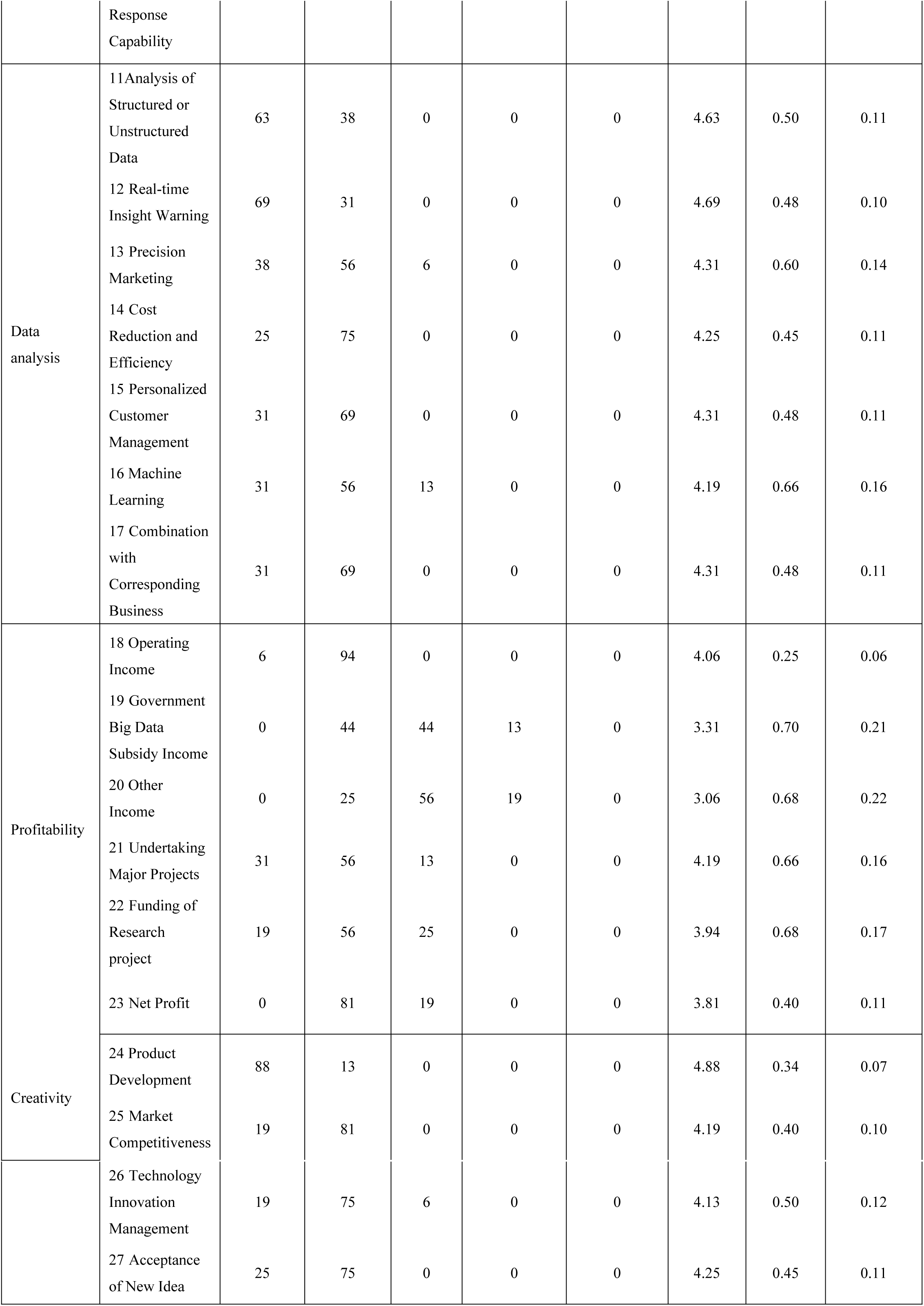
Importance for the first-level indicators in the second round

**Table 7.**
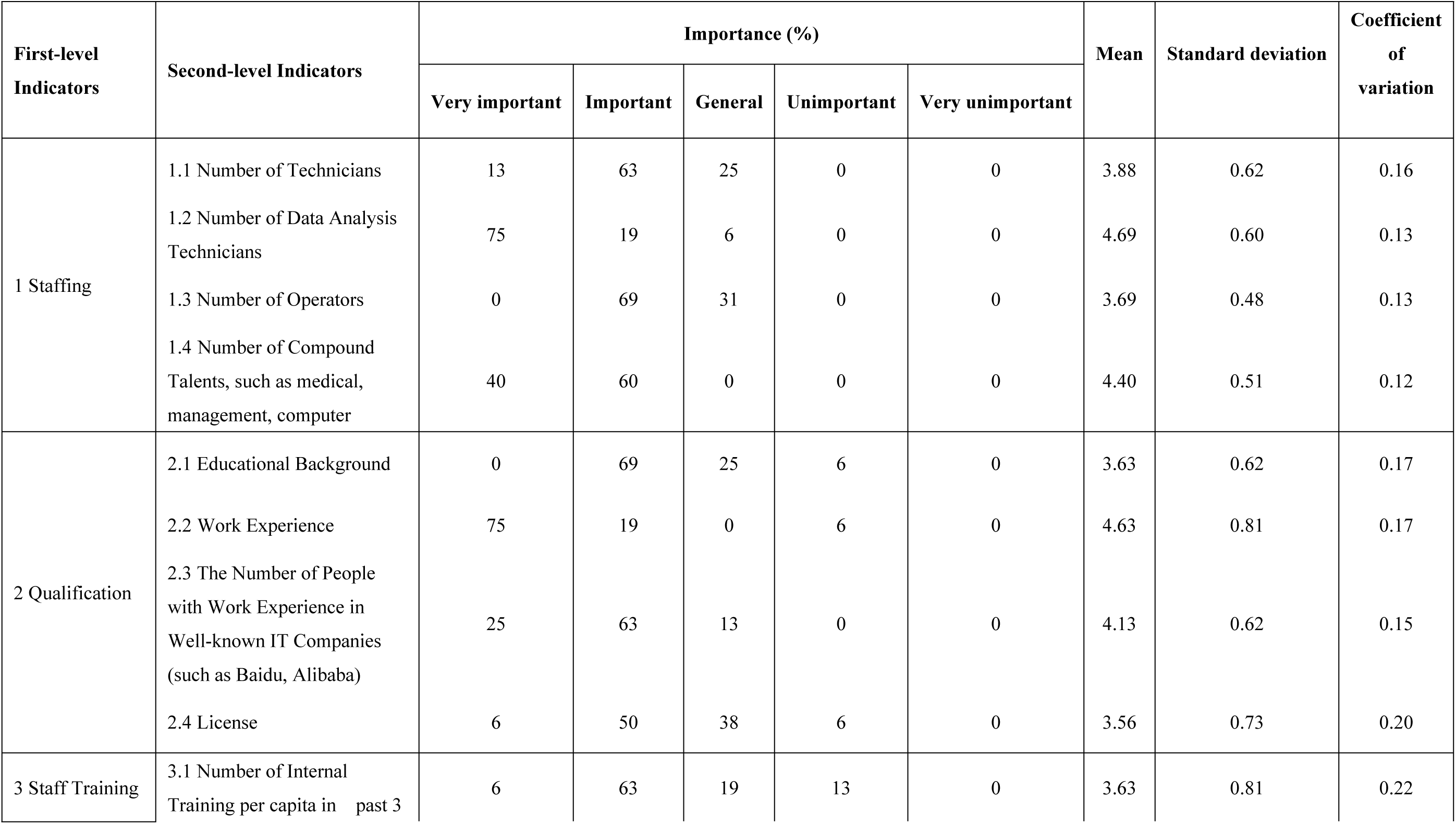
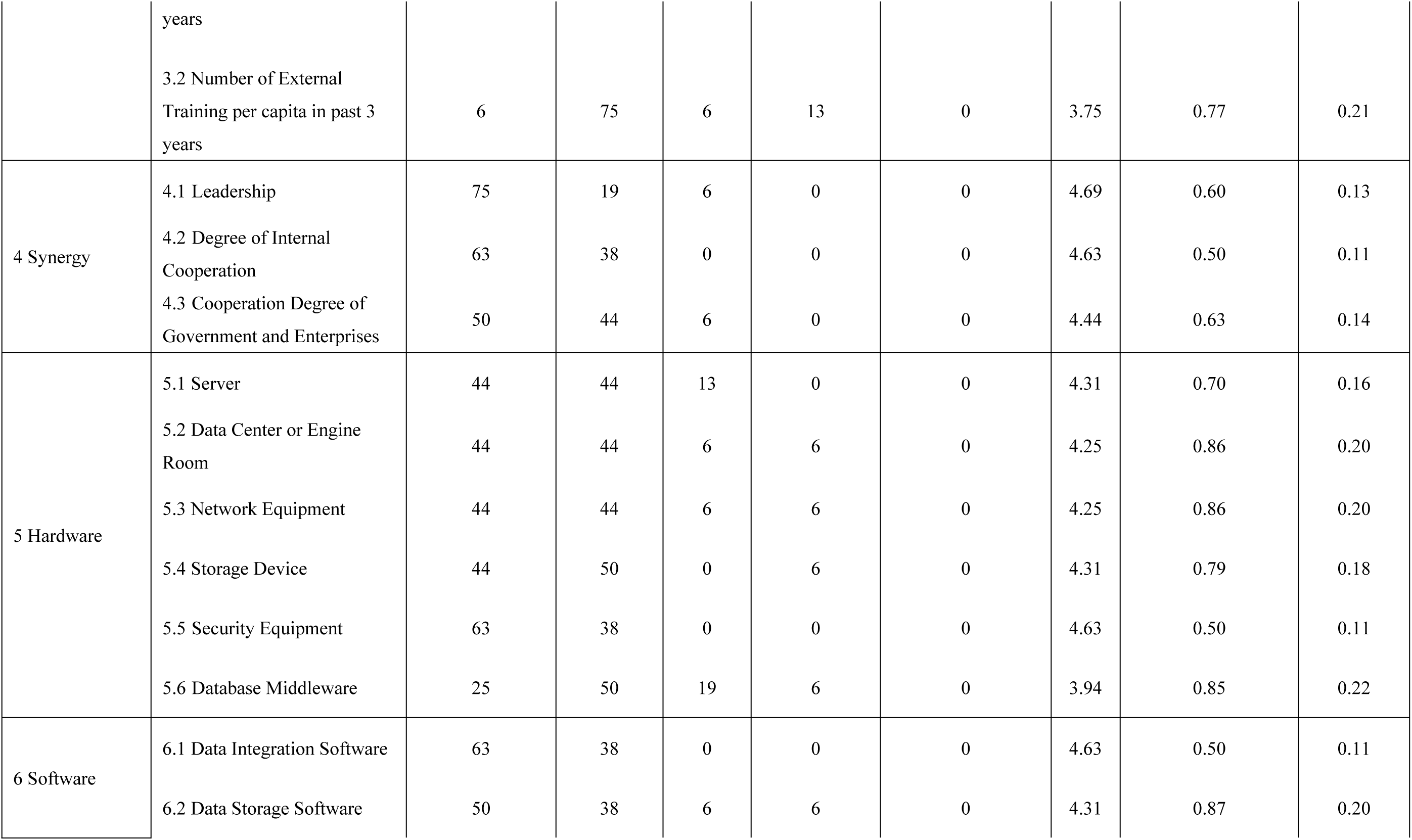
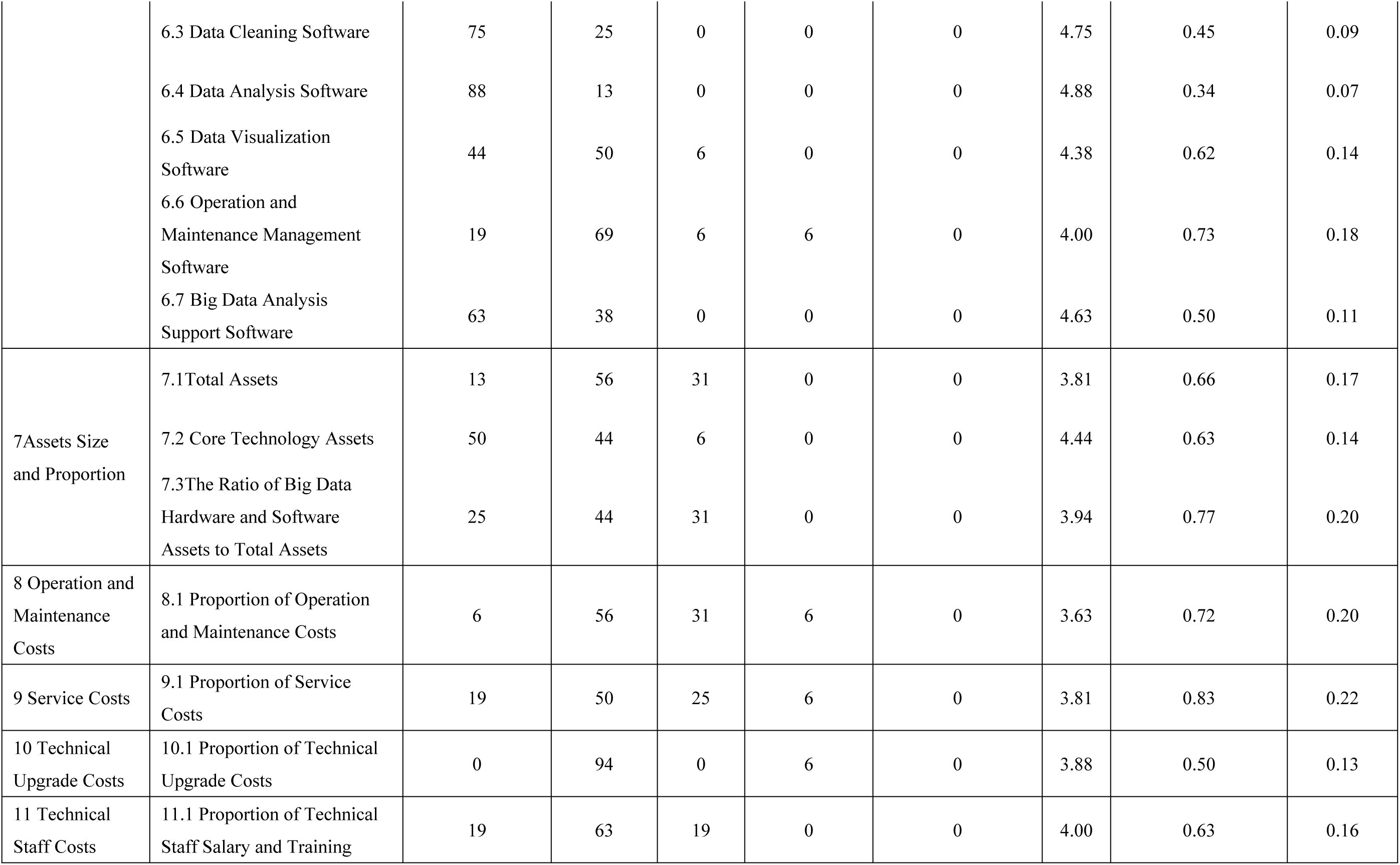
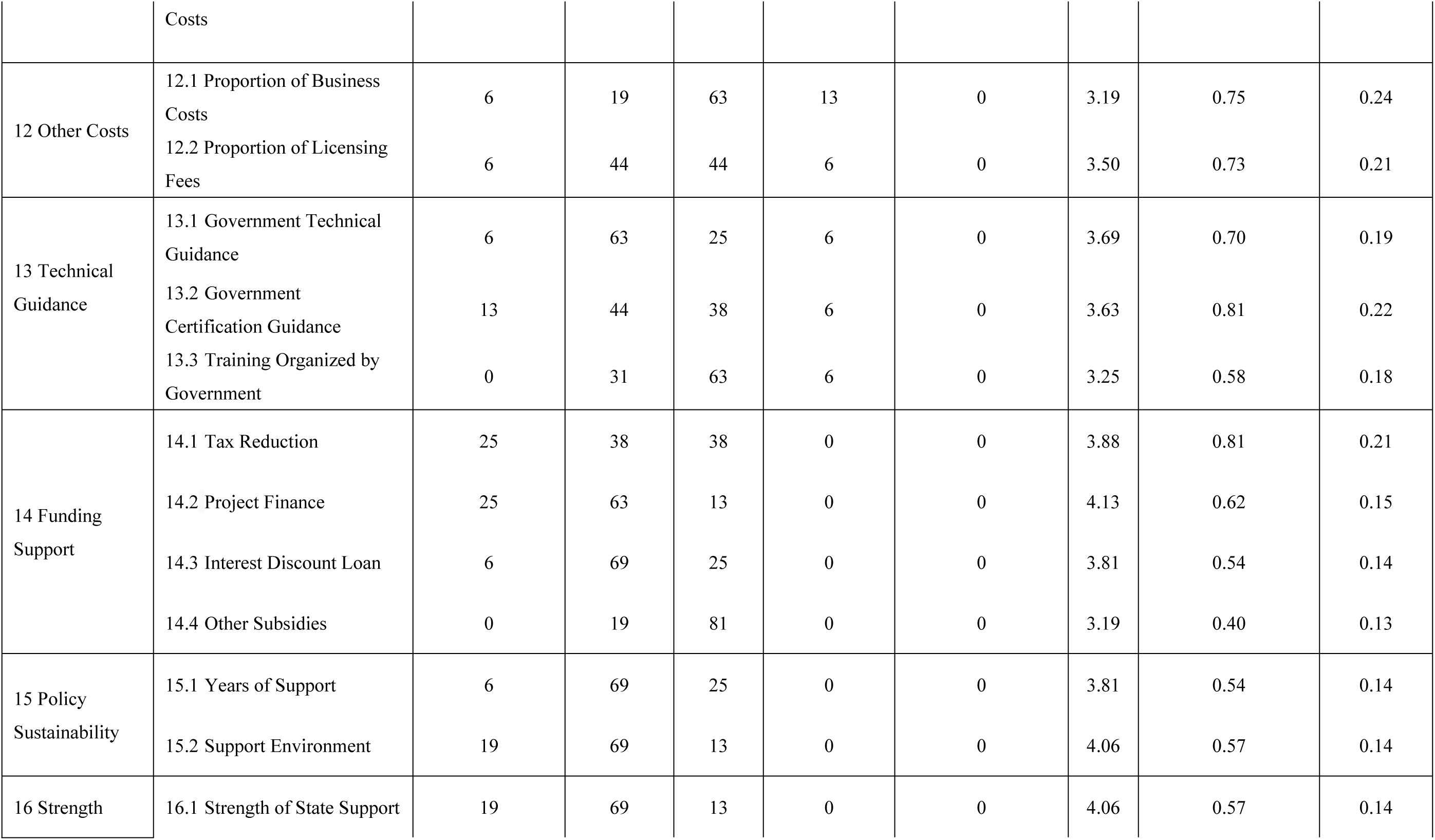
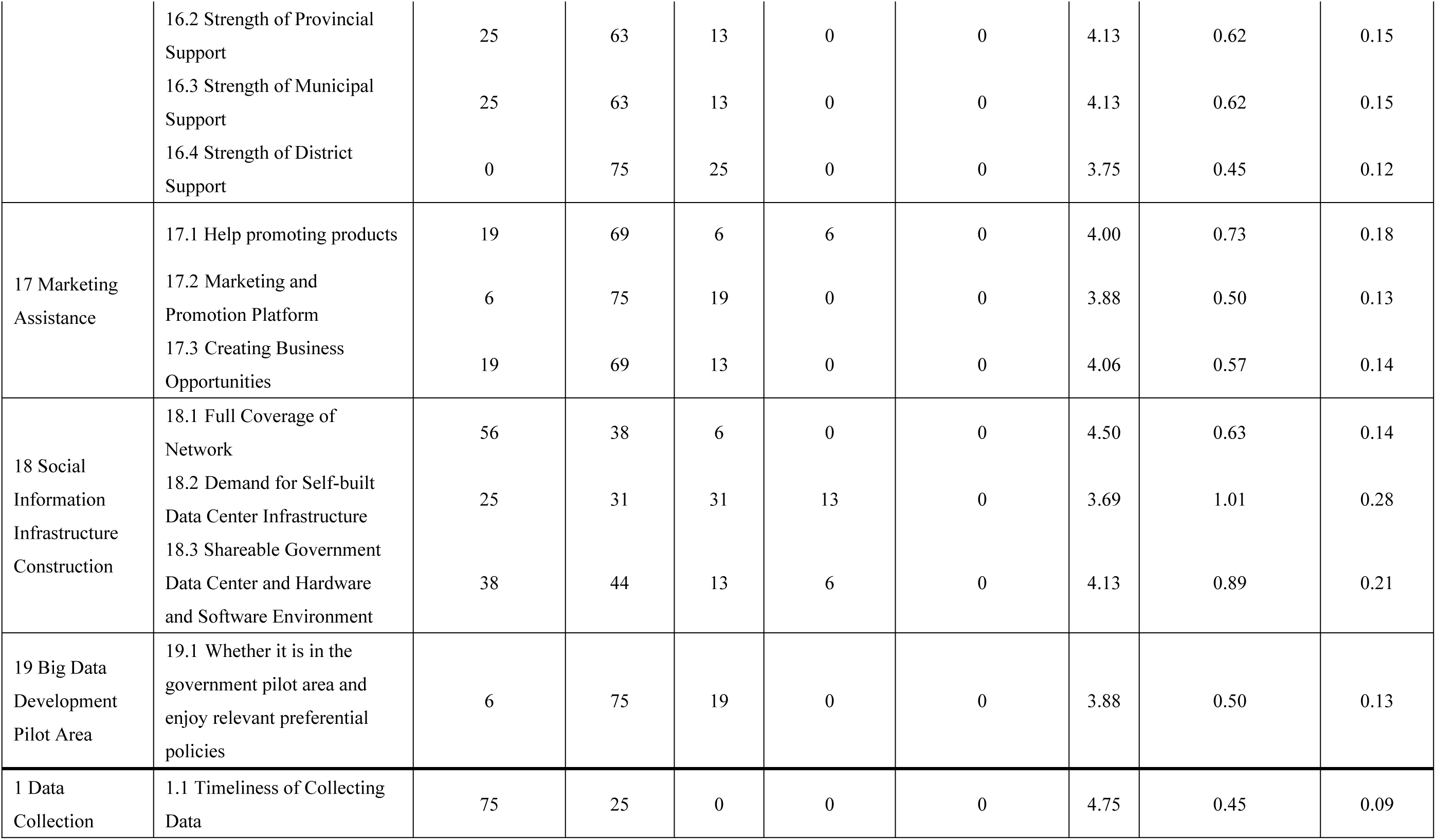
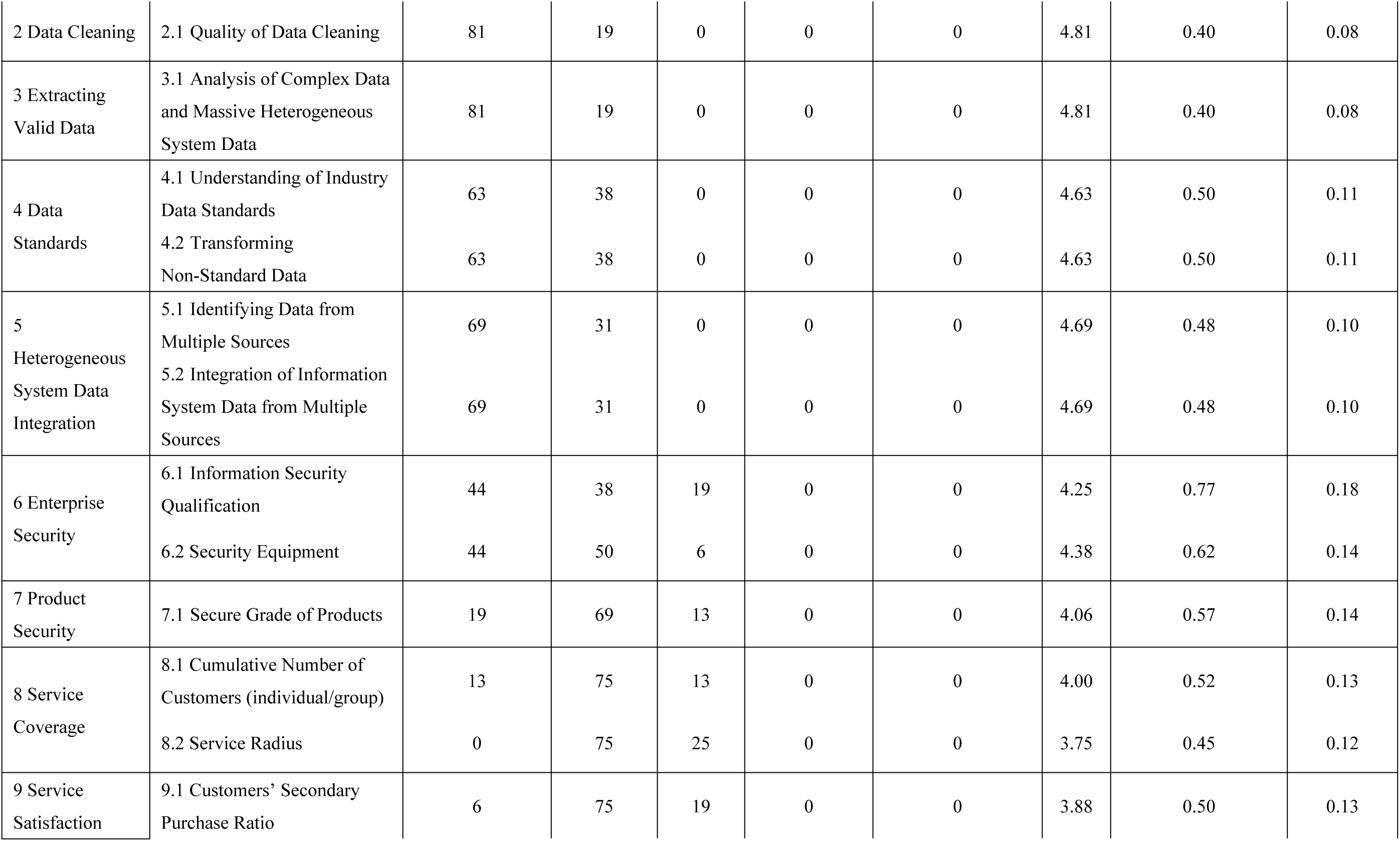
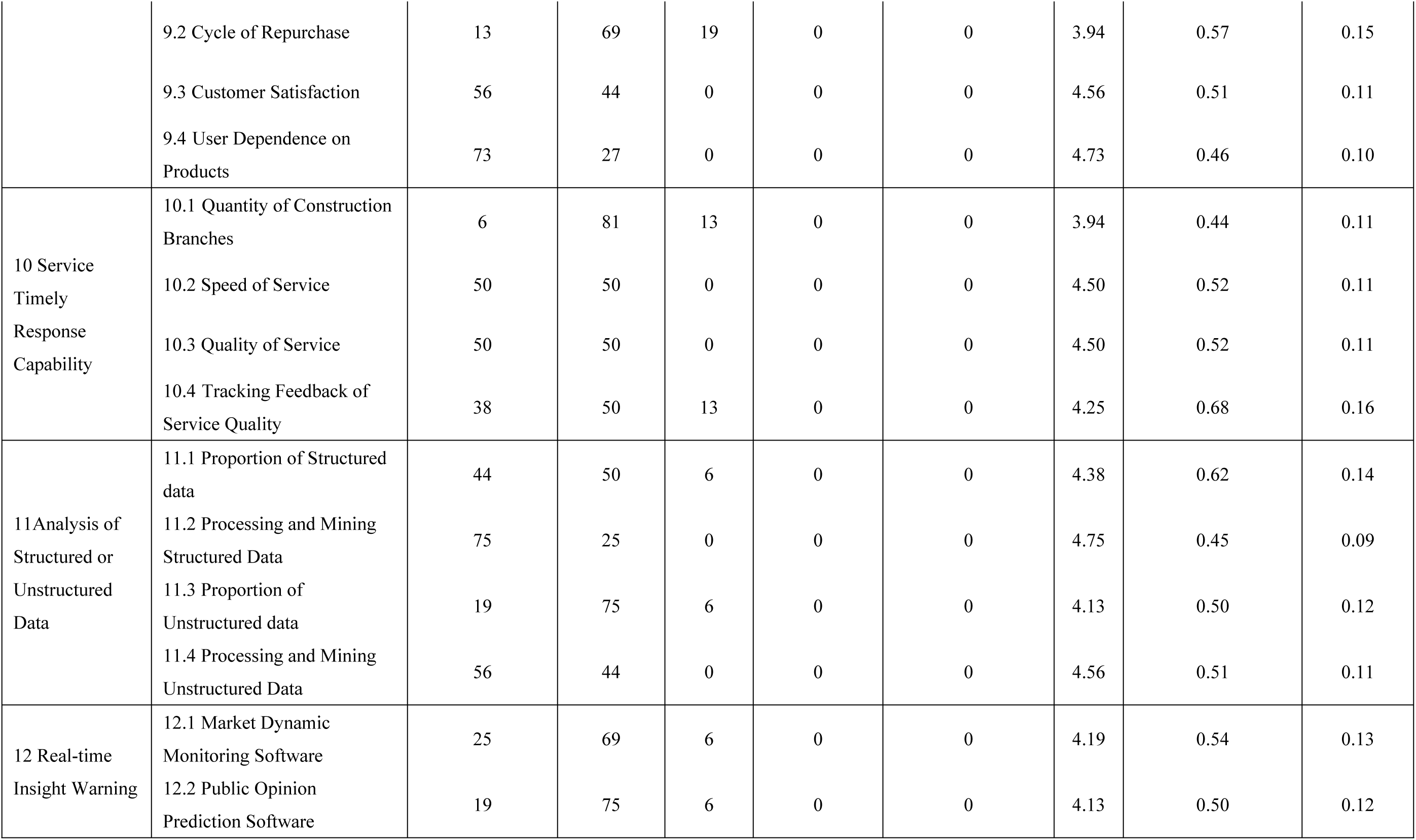
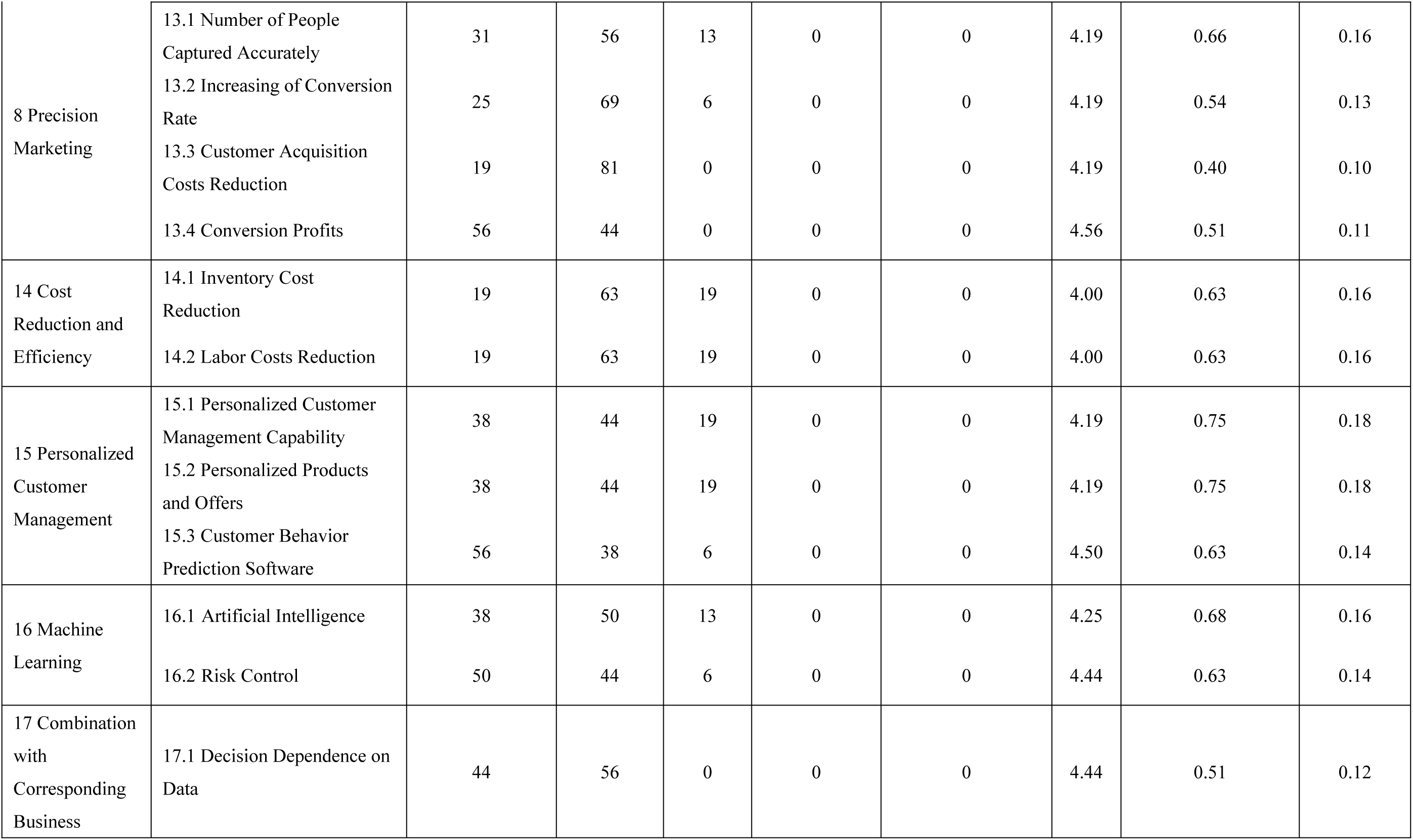
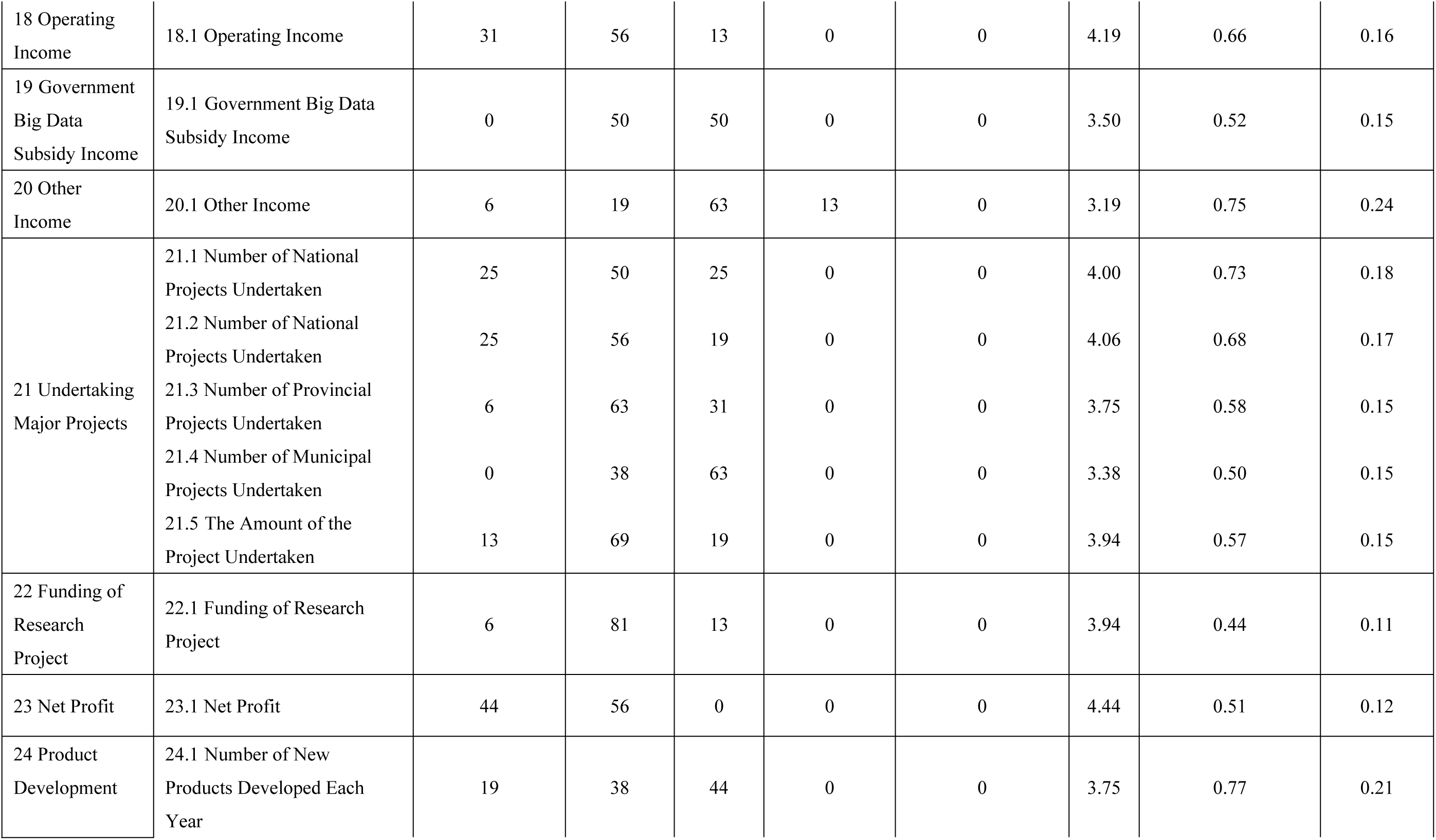
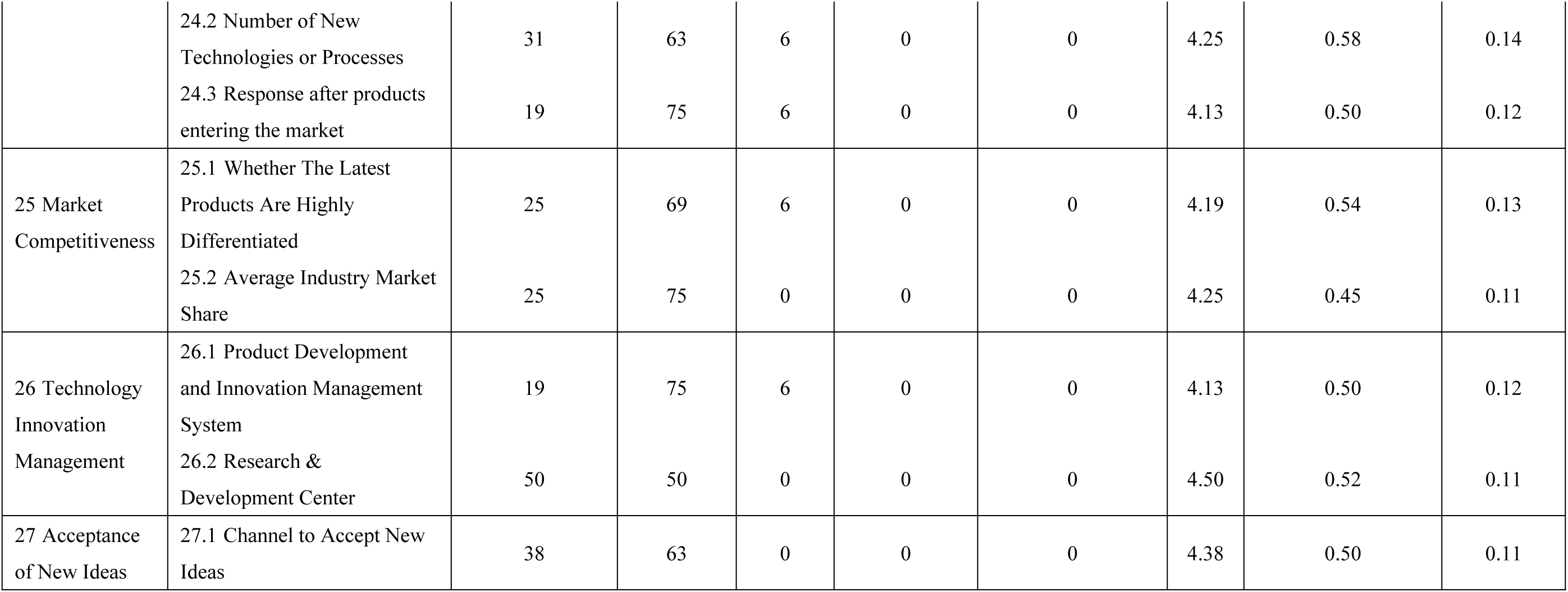
Importance for the second-level indicators in second round

After the second round, the research team again used the three statistical measures (importance percentage, mean score, and coefficient of variation) to evaluate the indicators. No indicator met all three conditions for deletion and the research team therefore did not delete any indicator.

From the second round of review, the dimensional consensus rate was 100%, the first-level indicator consensus rate was 87%, the second-level indicator consensus rate was 81%, and the consensus rate of all indicators was 84%, indicating that experts had high consistency in evaluation of the index. According to the Delphi method, the questionnaire could therefore be accepted.

## Conclusions

Based on the Delphi method, two rounds of expert opinions were used to develop the index system, and a health care enterprise big data application capability index system covering 11 dimensions, 46 first-level indicators and 111 second-level indicators was constructed.

The index system derived from this study can be used to evaluate the big data application capability of a health care enterprise. However, the basis of big data applications is all data rather than sample data. It is difficult for health care companies to grasp all the characteristics of customers. Data sharing between medical institutions, public health agencies, and health care companies can truly reflect user profiles. This requires an integrated and open information exchange platform between the company and the government. In the future, connectivity and interaction are the trajectories of big data applications. This index system focused on health care enterprise and lays the foundation for evaluating enterprises’ big data application. It is hoped that future research will deepen connections and interactive content of big data.

This study may be limited by research conditions and expert resources. This newly developed index system attempted to appraise the application capability of big data scientifically and systematically.

## References

1. Qingzhen, Liu., Transformation of enterprise marketing thinking under the new paradigm of "Internet +" technology and economy[J]. Academic Exchange, 2017. 1(1): p. 123–127

2. Li, Liu., On the Traditional View of Enterprise Benefits and Sustainable Development [J]. Economics and Management, 2004. 18(2): p. 34–36.

3. Ping, Zhang., Evaluation of Rural Health Service Capability — Taking J Province as an example [D]. PhD thesis of Jilin University, 2014.

4. Zhengguang, Niu., Research on the Impact of Big Data on the Modernization of Government Governance — Taking Hebi City as an Example [D]. Doctoral thesis of China Agricultural University, 2017.

5. Gupta Manjul, Gorge Joey F., Toward the development of a big data analytics capability [J]. Information & Management. Information & Management, 2016. 53(8): p. 1049–1064.

6. Wei, Jin., Report on the Competitiveness of Chinese Enterprises[R]. Beijing: Social Sciences Academic Press, 2003.

7. Xiaocong, Qian., The development of big data and industrial opportunities [J]. Internet of Things technology, 2013. 10: p. 84–86.

8. Yanling, Ma., The Study on Sustained Competitive Advantage of the Enterprise [D]. Master Degree thesis, Jilin University, 2004.

9. Chenyi, Qu., Cyberspace security and secrecy confrontation situation and coping strategies[J]. Confidential Science and Technology, 2012. 4: p. 6–11.

